# Infants who develop BPD have an airway endotype defined by vimentin expression and ciliary loss

**DOI:** 10.1101/2025.09.16.676660

**Authors:** Laurie C. Eldredge, Yan Han, Gail H. Deutsch, Jennifer M.S. Sucre, Shawyon P. Shirazi, Caroline Stefani, Stephan Pribitzer, Stephen R. Reeves, Lucille M. Rich, Elizabeth R. Vanderwall, Steven F. Ziegler, Jason S. Debley

## Abstract

**Rationale:** Bronchopulmonary Dysplasia (BPD) results from abnormal lung development after preterm birth, with structural deficits at every respiratory tree level. BPD with lower airway disease is emerging as a clinically significant phenotype with increased mortality, and there is a significant knowledge gap in the molecular mechanisms whereby preterm birth disrupts normal airway development.

**Objectives:** To develop a human model of lower airway disease after preterm birth and to characterize a molecular endotype of evolving BPD (eBPD) at baseline and in response to injury.

**Methods:** We used a combination of an *ex vivo* organotypic Airway Epithelial Cell (AEC models) and well-characterized pathologic and transcriptomic patient samples for quantitative immunohistochemistry and RNA sequencing analyses.

**Measurements and Main Results:** Compared to AECs from healthy patients, eBPD- derived AECs have a molecular endotype of reduced proliferation, impaired differentiation to ciliated epithelium, and an expanded vimentin-positive population with a transcriptional shift toward stromal cell-associated genes. With hyperoxia exposure, eBPD-derived AECs exhibited a pronounced vimentin response *ex vivo*, which parallels the increased vimentin expression of airway cells observed in lung tissue from human infants with BPD.

**Conclusions:** In this organotypic model of neonatal airway differentiation, we find that infants with eBPD have impaired differentiation, increased expression of vimentin, and concomitant loss of cilia, with an exaggerated increase in vimentin expression after hyperoxia injury, findings that mimic the effects of prematurity in airway cells in human patients. These data provide a foundation for future mechanistic studies interrogating the role of intermediate filaments in epithelial differentiation and repair.

## Introduction

Bronchopulmonary Dysplasia (BPD) is a severe chronic lung disease of infants born extremely premature, characterized by aberrant postnatal development and superimposed injury of the respiratory tree (1, 2). Historically, BPD research has focused on the aspects of the disease with arrest of alveologenesis, with the airway pathology being relatively understudied and underrepresented in animal models of BPD. BPD diagnostic criteria stratify the disease by clinical severity, and patients requiring invasive respiratory support at 36 weeks corrected gestational age (CGA) are diagnosed with Severe BPD (sBPD) Type 2 or Grade 3 BPD (2, 3). However, this classification does not fully capture the anatomical heterogeneity of the disease. While arrested alveologenesis is a hallmark pathologic feature of BPD, lower airway disease is also prevalent, with up to 90% of patients with sBPD having some degree of obstructive lung disease on infant pulmonary function testing (4). In addition, many patients have multilevel airway obstruction with gas-trapping and need for prolonged positive pressure support to assist with patent airways during exhalation; indeed, severe airway obstruction remains a relatively understudied aspect of BPD pathology. Here, we seek to tackle the knowledge gap in understanding the molecular and cellular factors associated with airway disease in patients with sBPD (4).

The airway epithelium serves many roles in the lung, including mucociliary clearance, barrier functions, and innate immune responses to pathogens. A well-differentiated pseudostratified airway epithelium in the airway may have more than 15 distinct cell types as identified across multiple existing lung cell atlases, including ciliated, goblet, and secretory cells that arise from (multiple subtypes of) progenitor basal cells (5–8). BPD and BPD-related airway epithelia have been underrepresented in most published tissue and single-cell analyses and while apical cilia, vital to movement of airway secretions and entrapment of pathogens prior to cough (9), have been shown to be structurally abnormal and dysfunctional in premature infants with evolving BPD (10), the mechanisms whereby abnormalities in airway epithelium relate to clinical respiratory disease in BPD have not been explored.

To this end, we used minimally invasive tracheal aspirates to characterize lower airway epithelial cells from patients with evolving BPD under normal and hyperoxia-exposed conditions to mimic the exposures of prematurity. We found abnormal structure and function in BPD-derived AECs and a failure of the normal airway differentiation program that associates with vimentin expression under organotypic culture conditions and in single-cell analysis of infant lung tissue.

## Methods

Detailed methods are provided in the data supplement.

### Airway Epithelial Cell Collection and Isolation

AECs were isolated from tracheal aspirates (TAs) collected as part of the routine respiratory care of endotracheally intubated premature infants born = <28 weeks gestation and who were less than 30 days of age in the University of Washington NICU (total N=27). BPD diagnosis and severity was assigned by LCE at 36 weeks CGA after chart review, per 2001 NIH guidelines and Abman et al., 2017 (11). Control AECs were derived from airway brushings collected from N=22 healthy children aged 2-16 years intubated for elective surgeries at Seattle Children’s Hospital. Both cohorts were obtained after consents per Seattle Children’s Hospital IRB-approved protocols (LCE STUDY00000263 and JSD STUDY#00001596). Parents of subjects provided written consent and children over 7 years of age provided assent. Details about AEC isolation and culture are provided in the supplement.

### Airway Epithelial Cell Organotypic Culture

Isolated AECs were differentiated for three weeks at air-liquid interface (ALI) with PneumaCult-ALI media in the basolateral compartment, as previously described (12–15). Quality control was performed in both control and BPD AEC cohorts to ensure they produced an organotypic differentiated epithelial cell culture with mucociliary morphology and without leak indicative of epithelial barrier. TEER measurements were also used on occasion to confirm barrier function, with additional details in the supplement.

### Hyperoxia-induced Epithelial injury of Organotypic AEC Cultures

Fully differentiated organotypic AEC cultures were exposed to 0.85 FiO2 for 24, 48, 72, or 96 hours in a hyperoxia chamber placed in the incubator (BioSpherix, Parish, NY), with 0.21 FiO2 exposed cultures as controls. Hyperoxia-induced epithelial injury was evaluated by visual inspection of cell morphology, immunostaining, and RNA sequencing. Increased expression of *GPX2* (16) and *NQO1* (17, 18) in the FiO2 0.85- exposed group was used to validate hyperoxia exposure through known responses to oxidative stress.

### RNA Sequencing and Analysis

Bulk RNA sequencing (RNA-seq) was performed on organotypic AEC cultures, with support from the Benaroya Research Institute Genomics and Bioinformatics Cores, with additional details in the supplement. RNA-seq datasets presented in this article will be submitted to the National Center for Biotechnology Information Gene Expression Omnibus by the time of publication.

### Immunostaining and Confocal Imaging

Proliferative or fully differentiated AEC cultures were washed with PBS, fixed with 10% formalin for one hour at room temperature, and labeled by wholemount immunofluorescence staining prior to quantitative confocal imaging, with additional details in the supplement.

### Analysis of Human Pathological Specimens

Pathological specimens from premature infants born at less than 26 weeks gestation who died from pulmonary/evolving BPD (N=4) or non-pulmonary (N=4) causes, were analyzed (IRB-approved protocol (GHD, Study00003664; Figure 7A). Those patients in the group noted to have non-pulmonary deaths were supported with oxygen and/or noninvasive respiratory support at birth/ DOL1 with milder lung disease, whereas those pulmonary causes of death required invasive respiratory support from birth/DOL1 on and had severe lung disease on pathological analysis. Formalin-fixed paraffin embedded 5-µm serial sections of trachea taken at the level of the thyroid were stained after antigen retrieval with anti-tubulin (Sigma, 1:10,000), anti- pancytokeratin (Dako, 1:100), anti-EPCAM (Invitrogen, 1:800), anti-vimentin (Biogenic, 1:100), and anti-TP63 (Biocare, 1:200). Staining was quantified in a blinded manner in three randomly selected areas of tracheal epithelium at 20X magnification and captured with a digital camera mounted on a Nikon Eclipse 80i microscope using NIS-Elements Advanced Research Software v4.60. Care was taken to avoid the denuded areas near the endotracheal tube.

### Data Analysis

Analysis of quantitative immunohistochemistry and total cell numbers was conducted with GraphPad Prism 8 (GraphPad Software Inc, La Jolla CA). Mann-Whitney or Student’s t-test were performed when appropriate to compare two treatment groups, depending on distribution of the data. Asterisks indicate statistical significance, *p<0.05, **p < 0.005, ***p < 0.0005 and **** p<0.0001.

### Single-cell RNA Sequencing Analysis

We interrogated a recently released single cell RNA-seq (scRNA-seq) atlas of human infant lung injury (8) from infants who died with BPD/BPD+pulmonary hypertension (N=4) and term infant controls (N=2) to look for differentially expressed genes in AECs. Full code used in the analysis of these data is available at https://github.com/SucreLab/HumanNeonatal. A table with the clinical characteristics of infants whose lungs were sequenced is in the supplement for the manuscript. In addition, raw data are available on NCBI GEO database (accession # pending) and raw and processed data are available at the LungMAP Consortium and as an interactive data explorer at www.sucrelab.org/lungcells.

## Results

### Identification of Deficient AEC Growth and Differentiation in Evolving BPD

We established an *ex vivo* system from tracheal aspirate-derived AECs (Figure 1A) to examine AEC proliferation and differentiation in evolving BPD (eBPD). We defined eBPD as infants born premature (<28 weeks), less than 30d old, who were later diagnosed with BPD per the standard definitions. Despite identical seeding densities, eBPD AECs grew more slowly than healthy control AECs, and most eBPD AECs (7/12, 58%) had features of failed differentiation including lack of apical polarization/topography, evidence of impaired barrier function with apical leak, and altered cell morphology. By comparison, only 1/6 control AEC organotypic cultures had any features of poor differentiation. eBPD AECs stained for apical cilia and actin displayed disorganized morphology with stretched actin network, decreased ALI thickness, and scant apical cilia relative to controls (Figure 1B).

**Figure 1.**
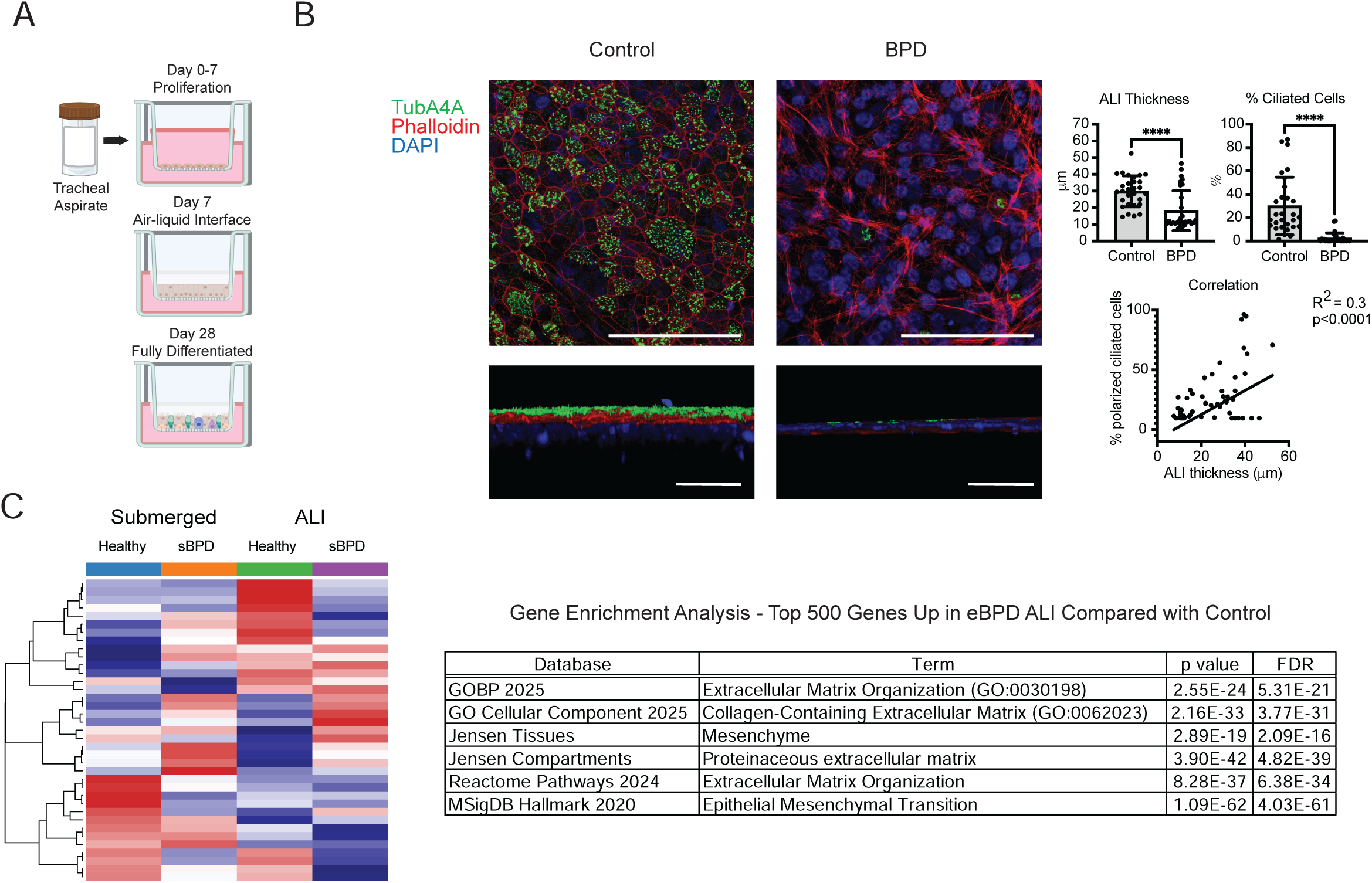
Establishment and RNA Sequencing of Organotypic Airway Epithelial Cell Cultures from eBPD Tracheal Aspirates. (A) Organotypic AEC cultures were generated from unmodified tracheal aspirates isolated from intubated premature infants, grown in submerged culture until confluent, and then differentiated at air-liquid interface (ALI). (B) 3D morphological assessment of AEC differentiation, which includes quantitative immunostaining of apical cilia, polarization, and ALI thickness. Decreased ALI thickness, % polarized ciliated epithelium, and correlation between these two measures, in demonstrated in organotypic cultures from eBPD patients (N=6), compared with controls (N=6). (C, left) Heat map with transcriptomes of healthy pediatric controls and eBPD patients in submerged and ALI conditions. (C, right) Gene Enrichment analysis of top 500 genes upregulated in eBPD and downregulated in control AECs after differentiation at ALI, demonstrating differential expression of genes controlling extracellular matrix across multiple databases with an emphasis on epithelial-mesenchymal transition in Hallmark Gene Analysis. (https://maayanlab.cloud/Enrichr). (Green = anti-TubA4A, Red = phalloidin, Blue = DAPI. ****p< 0.0001, Mann-Whitney test, magnification bar = 100 μm)

Bulk RNA-seq of control and eBPD AECs was performed in both proliferative (day 7 of submerged condition) and differentiated cultures. There were 5235 differentially expressed genes (DEG) between control proliferative and differentiated cultures, 979 DEG between eBPD proliferative and differentiated cultures, and 1842 DEGs between control and eBPD differentiated cultures (Figure 1C, Table 2). In Gene Enrichment analyses, the 500 most significantly upregulated genes in eBPD samples were associated with extracellular matrix (ECM) gene expression and epithelial mesenchymal transition (EMT), as detailed in Figure 1C. Expression of traditional hallmark genes of epithelial-mesenchymal transition was increased, including CDH2 (9- fold) and VIM (6-fold) in eBPD cultures in comparison with controls.

We next attempted to “rescue” eBPD cultures by using a more supportive medium that was becoming more widely used as these experiments evolved (see methods). With this new medium, the failure rate of differentiation in eBPD cultures decreased to 36% but still was greater than controls (<10%). Despite healthier organotypic AEC cultures with increased proportion of ciliated cells in both groups cultured in this medium, eBPD cultures had persistently decreased ALI thickness and apical cilia compared with control cultures (Figure E1).

We next compared the growth potentials of eBPD and control AECs by quantifying P63+ basal cells in proliferative cultures and total cell numbers in fully differentiated cultures. Proliferative AECs from eBPD samples had fewer P63+ basal cells at day one, three, and seven days after plating on transwells - and fewer cells overall in fully differentiated cultures - despite consistent plating density (Figure E2). Bulk RNA-seq comparing gene expression in control and eBPD AECs in proliferative (Supplementary Tables E1 and E2) and fully differentiated ALI cultures (Supplementary Table E3) was performed. TP63 and Ki67 expression were decreased in eBPD AECs from all timepoints, compared with controls.

Taken together, these data demonstrate that AECs from premature infants with eBPD do not differentiate normally and upregulate ECM gene expression modules that may be associated with extracellular remodeling and/or other changes in AEC function specific to BPD pathophysiology.

### Identification of Vimentin-expressing AECs in eBPD

Building on the finding of increased EMT-associated gene expression, we interrogated the mesenchymal transcriptional signature of cells in AEC cultures from patients with eBPD and healthy controls and found that AECs from patients with eBPD had increased expression of vimentin by quantitative immunostaining and bulk RNA-seq (Figure 2A). Vimentin-positive cells were present in similar numbers of control and eBPD AECs after 3 days of proliferation. In contrast, there was an increase in the number of vimentin-positive AECs in eBPD AECs after 7 days in proliferative conditions with a further increase in fully differentiated AECs, compared with controls (Figure 2B). Bulk RNA-seq in well-differentiated control (N=5) and eBPD (N=5) cultures demonstrated 22-fold greater vimentin expression in eBPD cultures (Figure 2C). We used several methods to distinguish poor AEC differentiation in eBPD associated with vimentin expression from stromal contamination of the seeding population of cells (Figure E3).

**Figure 2.**
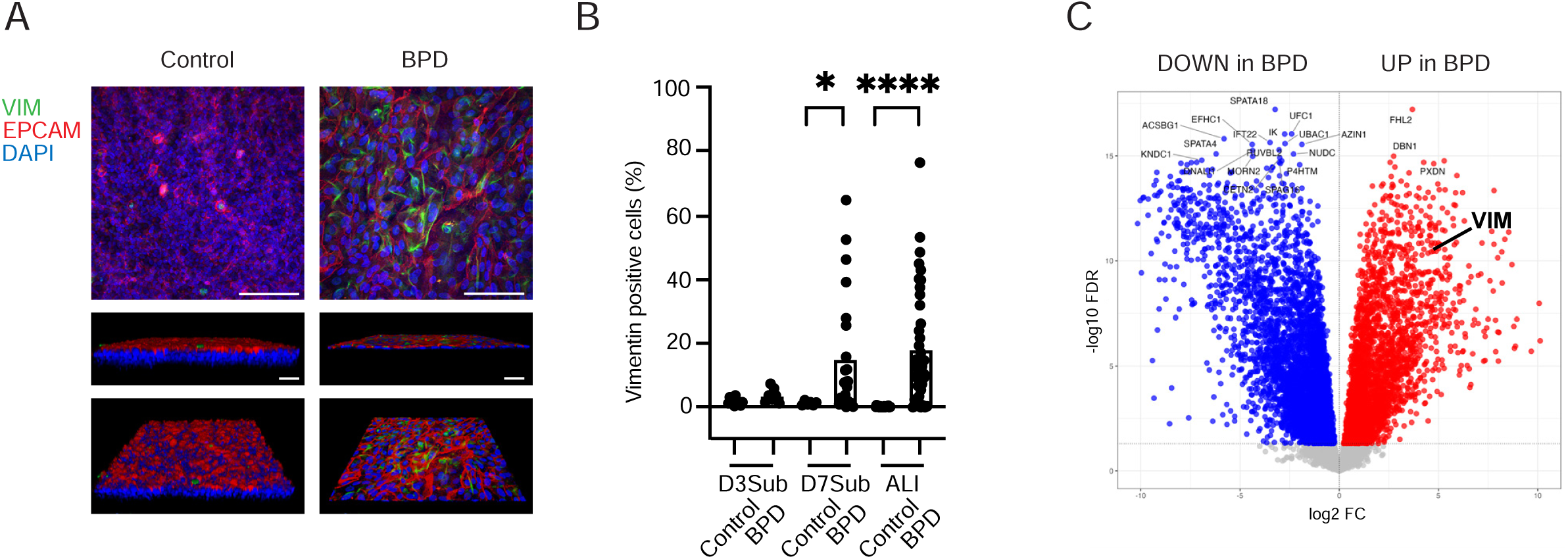
Increased Vimentin-expressing AECs from Patients with eBPD. (A) 3D projections of vimentin immunohistochemistry on organotypic AEC cultures from eBPD and control patients. (B) Quantification of VIM+ cells during proliferation phases (Day 3 and Day 7 submerged cultures) and at ALI after 3 weeks of differentiation, with increased VIM+ cells in eBPD samples at day 7 and at ALI. One dot = one image, with N=2-3 images per condition, from total of N=5 control donors at day 3, N=4 eBPD at day 3, N=3 controls at day 7, N=6 eBPD at day 7. (C) Bulk RNA Sequencing on N=5 Control and N=5 eBPD differentiated cultures, showing 22-fold upregulation of vimentin expression in eBPD AECs. (Green = anti-vimentin, Red = anti-EPCAM, Blue = DAPI. *p<0.05, ****p< 0.0001, Mann-Whitney test, magnification bar= 100 μm)

### Additive Increases in Vimentin Expression after Hyperoxic AEC Injury

We employed hyperoxia exposure to model the airway disease associated with BPD by treating fully differentiated AECs with 0.85 FiO2 or normoxia for 24, 48, 72, or 96 hours and investigating structural and molecular changes after hyperoxic injury. We found a dose-dependent response in loss of apical cilia, with increased ciliary loss after subsequent days of hyperoxia in control cultures (Figure 3B). At baseline (normoxia), there was an increased number of VIM+ AECs in eBPD cultures compared with controls (Figure 4A) that was consistent with mRNA expression differences (Figure 2C). Each day of hyperoxia exposure further increased the number of VIM+ AECs in both control and eBPD groups (Figure 4B).

**Figure 3.**
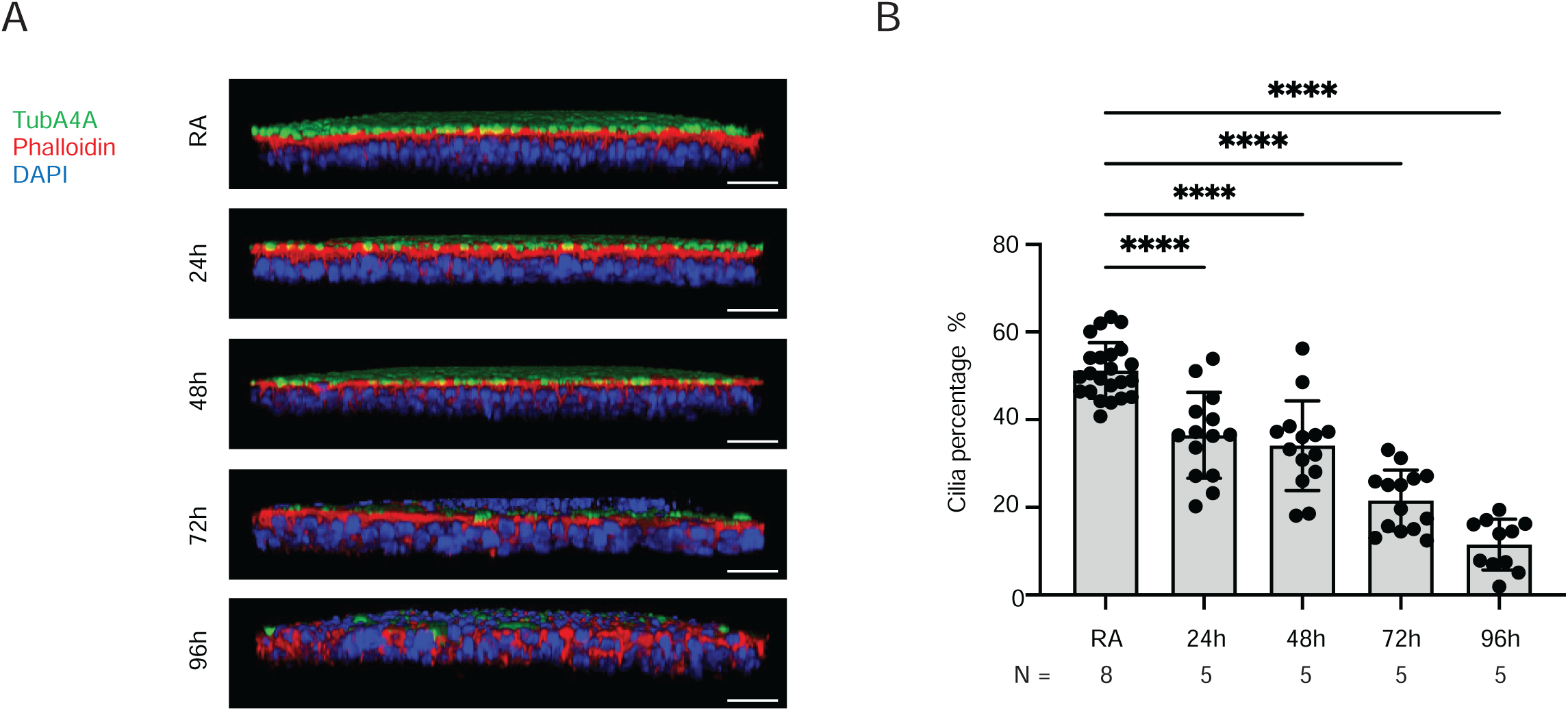
Hyperoxia-induced AEC Injury *Ex Vivo* With Loss of Cilia. (A) Immunostaining of organotypic AECs from healthy controls in room air (RA) or after exposure to hyperoxia for 24, 48, 72, or 96 hours. Increased loss of apical cilia (green) with additional days of exposure to 0.85 FiO2. (B) Hyperoxia-induced dose-dependent ciliary loss, with % of ciliated culture quantified in N=2-3 images per condition in N=4-8 donors per condition as specified. (Green = anti-TubA4A, Red = phalloidin, Blue = DAPI. ****p<0.0001, Mann-Whitney test, magnification bar = 100μm)

**Figure 4.**
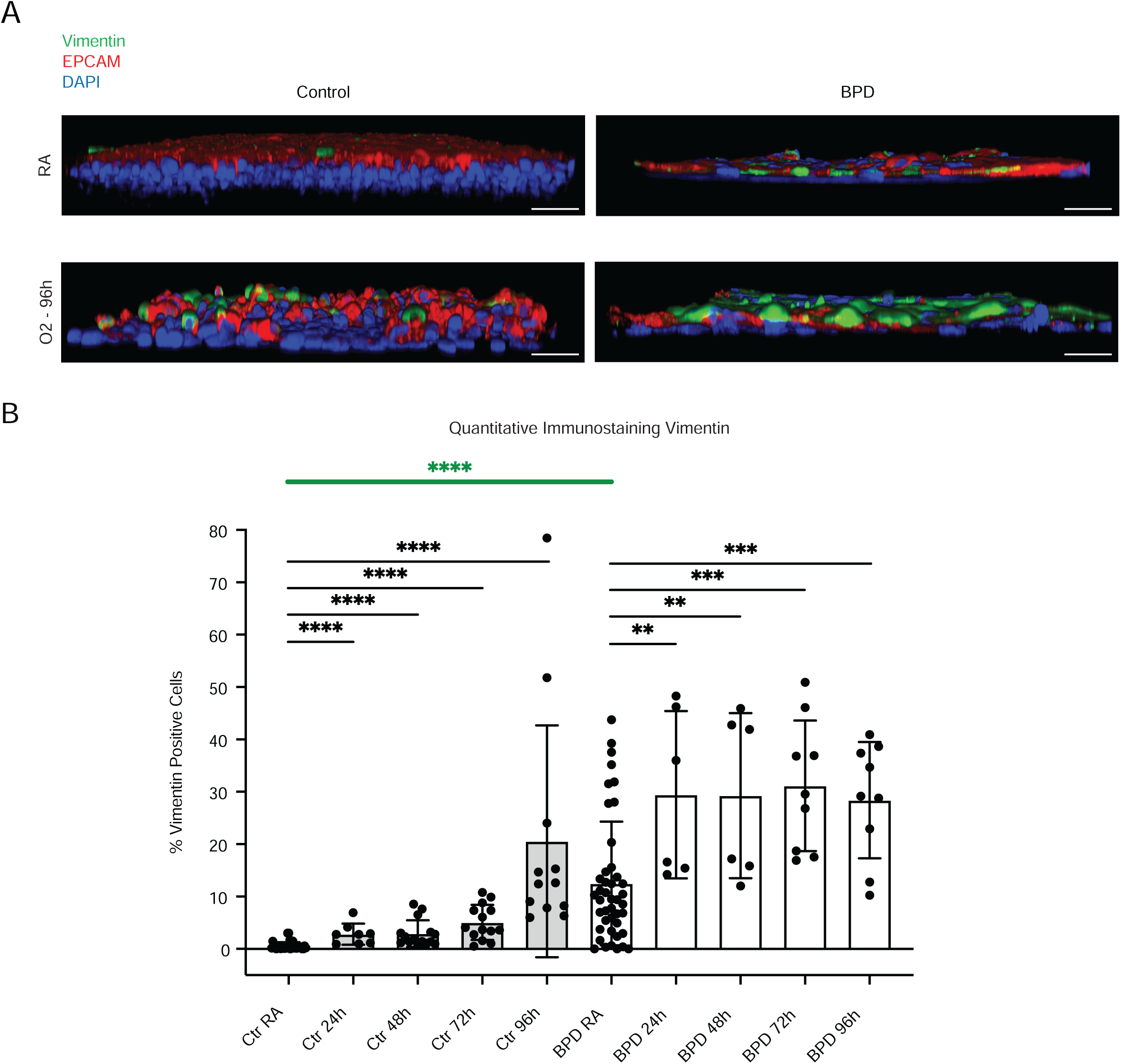
Increased VIM-expressing AECs in BPD at Baseline and After Hyperoxia-induced Epithelial Injury *Ex Vivo*. (A) Immunostaining of organotypic AEC showing increased vimentin expression (green) in BPD cultures at baseline (room air, 0.21 FiO2) and after 96h of exposure to 0.85 FiO2. (B) Quantification of VIM+ cells in room air or hyperoxia-exposed conditions, with comparisons of control and eBPD organotypic cultures as indicated. Green line indicates baseline (room air) differences of VIM+ cells between control and eBPD AECs. One dot = one image with N=2-3 images per N=3-8 donors per condition. (Green = anti-vimentin, Red = anti-EPCAM, Blue = DAPI. **p<0.005, ****p<0.0001, Mann-Whitney test, magnification bar = 100μm)

### Increased Vimentin Expression in BPD AECs at the Single Cell Level

We next explored cell-specific vimentin expression in BPD using a publicly available scRNA-seq atlas (8). The authors sequenced 43,607 single cells, and clustering analyses with cell-type annotation identified seven epithelial clusters from N=2 term controls and N=4 BPD samples isolated from human lung tissue after autopsy (Figure 5A). Five distal lung AEC populations were identified, including basal, multiciliated, RASC, Secretory SCGB3A1/SCGB3A2, and Secretory MUC5B cells (Figure 5B). Vimentin was localized to RASC and Secretory SCGB3A1/SCGB3A2 AECs, and vimentin expression was increased in these subpopulations in BPD samples compared with term controls (Figure 5C). In the 763 non-ciliated respiratory epithelial cells, but not other airway epithelial subtypes, vimentin expression was increased in the BPD samples, compared with unchanged expression of EPCAM between control and BPD cells (Figure 5D), supporting the possibility that increased vimentin expression is linked to airway injury and maladaptive repair after preterm birth. Interestingly, vimentin expression in BPD mesenchymal cells is also slightly increased compared with control mesenchyme (data not shown).

**Figure 5.**
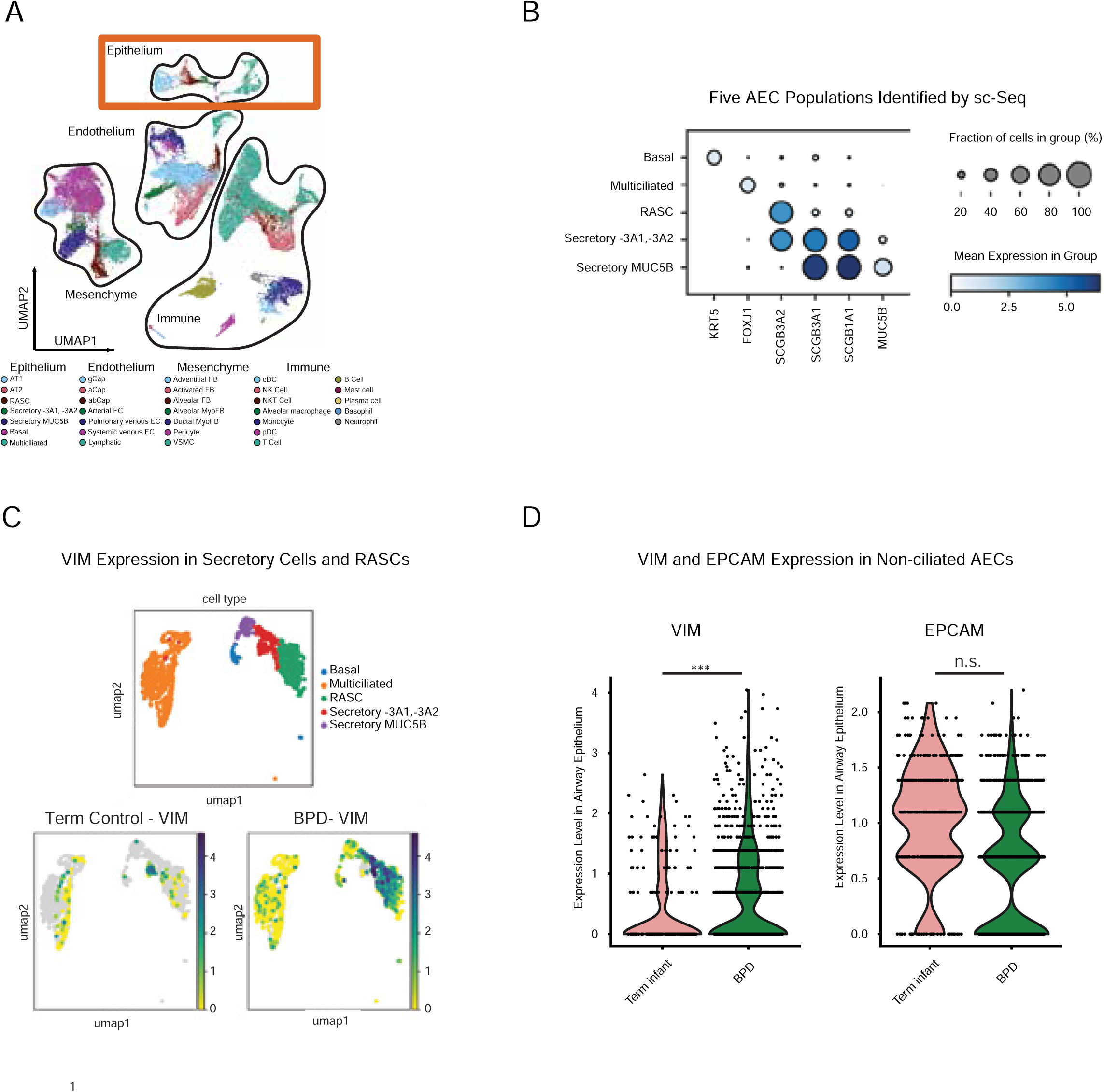
Increased Vimentin Expression in Non-ciliated, Secretory AECs Isolated from Human Lung tissue from Patients with Established BPD. (A) Single cell RNA Sequencing library created from lungs of patients with BPD (N=4) compared with term controls (N=2). (B) Five distal lung AEC populations were identified by cell-specific markers, including basal, multiciliated, RASC, and Secretory SCGB3A1/SCGB3A2, and Secretory MUC5B cells. (C) Localization of vimentin expression to RASC and Secretory SCGB3A1/SCGB3A2 AECs in BPD samples. (D) Vimentin, but not EPCAM, expression was increased in 763 non-ciliated airway epithelial cells isolated from pathological specimens from patients with BPD compared with term controls. (***p<0.0005 Mann- Whitney test).

### Aberrant AEC Differentiation and Increased Vimentin Expression in Pathologic Samples from Patients with eBPD

We sought to confirm our findings by investigating tracheal epithelial cells in pathologic samples from premature infants with and without eBPD. Controls were born extremely premature, did not require intubation on DOL1, had lower oxygen requirements in the 48-72h prior to demise, and had minimal lung disease on pathology (blinded analysis by GHD, Figure 6). Patients with eBPD were born extremely premature, required invasive mechanical ventilation on DOL1, had high oxygen requirements prior to demise, and had confirmed lung pathology including patchy lobular hyperinflation, airway mucostasis and hypertensive changes of the pulmonary arteries. Concordant with airway mucostasis in the evolving BPD cases, tracheal epithelia had severely reduced area of apical cilia compared with age-matched premature infants (***p<0.0005, Figure 6B). Additionally, we confirmed increased vimentin expression in tracheal AECs from patients with evolving BPD (*p<0.05).

**Figure 6.**
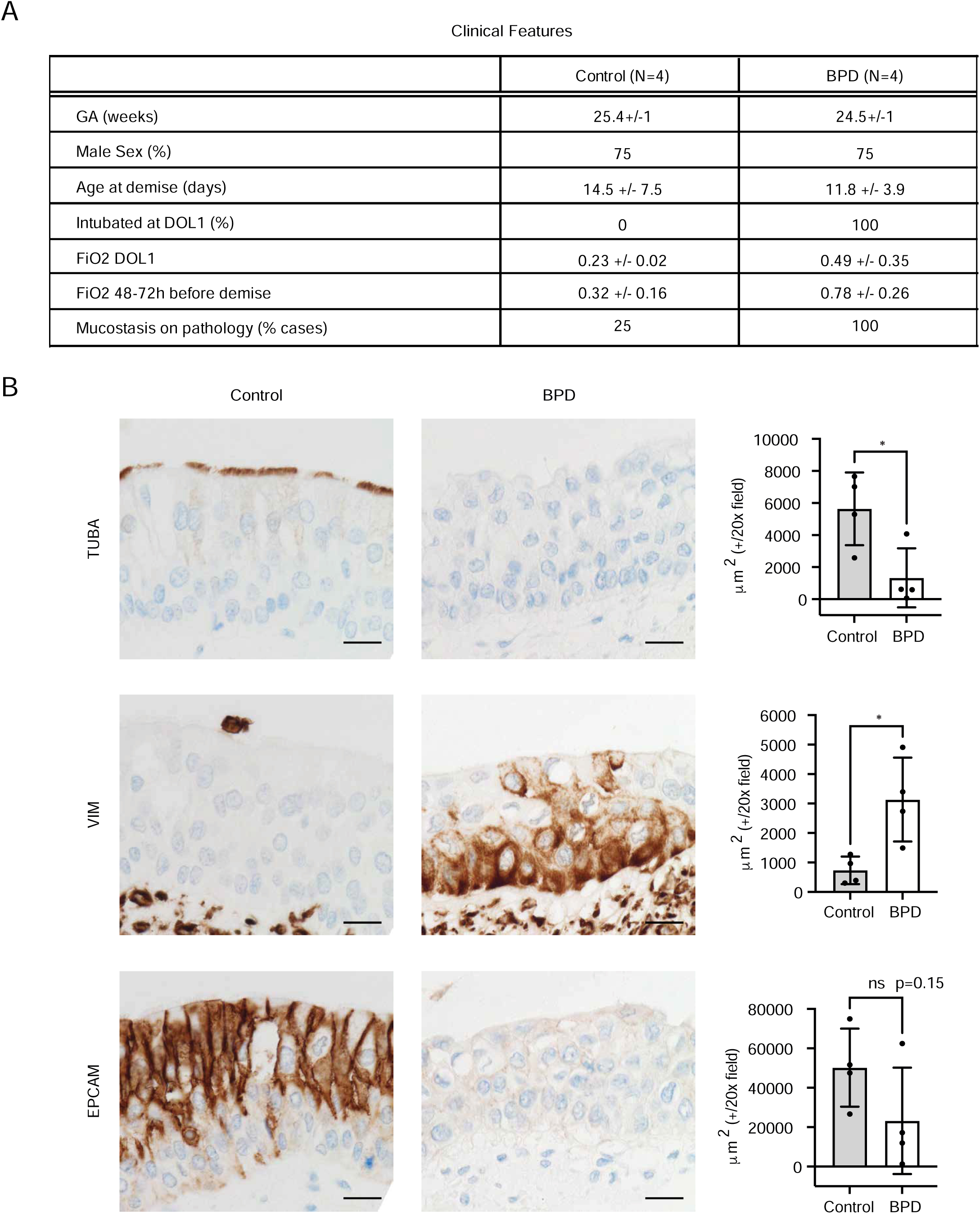
Correlating AEC Differentiation Defects *Ex vivo* with Complementary Analyses in Pathologic Samples from Patients with eBPD. (A) Clinical data from premature control (N=4) and eBPD (N=4) patients whose autopsies were reviewed. Control patients were born premature but supported with non-invasive respiratory support at birth and had milder lung injury on pathological analyses. eBPD patients required invasive respiratory support at birth with higher FiO2 and had severe lung injury on pathology. Tracheal AECs from premature infants (GA 23-26 weeks) who died from eBPD have decreased apical cilia (stained with anti-TUBA4+), increased VIM+ cells, and decreased expression of the cell adhesion marker EPCAM, compared with premature controls who died from non-pulmonary causes. (Values are mean +/- SD, *p<0.05, Mann-Whitney test, magnification bar = 20μm).

Tracheal AECs from patients with evolving BPD also had a trend toward decreased EPCAM (p=0.15) and increased basal cell TP63 (*p<0.05) expression.

## Discussion

In a combination of orthogonal techniques with primary neonatal human data, we demonstrate that AECs from premature infants who develop severe BPD exhibit deficient growth and differentiation when compared with AECs from pediatric healthy controls. As sampling the lower airway in premature infants is inherently challenging, we developed an *ex vivo* organotypic AEC model and demonstrated that AECs from patients with evolving sBPD fail to undergo the normal differentiation program to the polarized, pseudostratified respiratory epithelium observed in AECs from healthy pediatric donors. Further, these sBPD AEC cultures expressed more vimentin RNA and protein. When modeling the hyperoxia-induced injury associated with preterm birth by exposure of a subset of *ex vivo* cultures to 0.85 FiO2, we observed an increase in non-ciliated AEC-specific expression of vimentin in both control and sBPD AECs, a finding we confirmed in postmortem infant airways and a recently generated single cell atlas of neonatal lung injury. Together, these data highlight a new role for vimentin in evolving airway injury in preterm infants that correlates with response to hyperoxia and risk of BPD severity.

There is a growing appreciation for the different anatomic manifestations of BPD, and many survivors of prematurity have life-long lower airway disease (19–23). Small airway obstruction in these patients is multifactorial, a convergence of congenitally narrowed airways, dysanaptic growth, and repeated airway injury and repair. How the airways of patients born prematurely grow and remodel in the neonatal period is not well understood due to an absence of airway function data in during the neonatal period.

Moreover, there are not good predictors for which infants with evolving BPD will develop lower airway obstruction and hyperinflation. Modeling airway epithelial differentiation using human samples from well characterized donors with and without BPD has the potential to greatly advance understanding of mechanisms driving AEC growth, differentiation, repair, and function in BPD and identify potential novel therapeutic targets. In contrast to recent reports deriving airway basal stem cells from human pluripotent stem cells (iPSCs)(24) and basal stem cells from tracheal aspirates using EpCAM+ selection and/or mTOR inhibition (25, 26), our organotypic AEC model derives cells from unmodified tracheal aspirates. A strength of this approach is the preservation of the cellular diversity in the airway niche, replicating more of the cellular and molecular deficits that accompany eBPD. In our *ex vivo* model of AEC growth and differentiation, we identified a primary differentiation defect in AECs from patients with eBPD. Also, organotypic AEC cultures from patients with evolving BPD demonstrated distinct transcriptomic signatures with increased expression of matrix associated genes, a finding replicated in tissue and single-cell transcriptomics from human patients. In addition to baseline differences, we observed marked changes in the hyperoxia response, with decreased apical cilia in AECs from BPD patients. This replicates a known marked loss of AEC apical cilia and mucociliary clearance in BPD (10), an underappreciated driver of airway disease in this population that is not faithfully mimicked in most animal models (27, 28). Data from our organotypic cultures suggest that the loss of ciliated epithelia dovetails with an apparent expansion of vimentin- expressing AECs; future work to discover how this reorganization of AEC subtypes affects airway structure and function and the lineage and spatial relationships of epithelial cells in the premature airway will be crucial to a mechanistic understanding of airway development and BPD.

Vimentin is an intermediate filament protein with a known role promoting alveolar wound repair in adult lungs and driving cellular migration in the wound scratch assay (29). From our data, we conclude that the AEC shift to vimentin-positive cells is an indicator of AEC plasticity, but whether it is both a driver or merely an indicator of injury remains undetermined. Vimentin is likely helpful and necessary for the remodeling and repair of injured lower airway epithelia, though potentially at the expense of mucociliary clearance. In alveolar epithelial repair, there is growing appreciation of alveolar type 2 cell expression of intermediate filament keratin-8 as both a marker and mediator of fibrosis after injury, with genetic knockout of keratin-8 preventing fibrosis in mouse models (30). Whether these changes are transient or sustained is an important question, especially in light of long-term airway obstruction in survivors of BPD (19–23). With larger patient cohorts, and a longitudinal analysis of cultures from patients with evolving and progressive BPD, we plan to determine the trajectory of airway epithelial changes – including vimentin and other mediators- in the continuum of injury to repair.

There are several limitations to this study. AECs from healthy children and patients with evolving BPD have different sources (bronchial brushings versus tracheal aspirates), but airway brushings were not feasible in the first 30d of life. Also, the number of AECs in tracheal aspirates was smaller and variable (200–1000), and therefore these cells likely underwent more doublings in establishing ALI cultures. To mitigate this, we prioritized complementary and concurrent analyses with pathological samples from patients with evolving BPD. In addition, as being intubated is required for obtaining a tracheal aspirate, our analyses include a subset of patients with more severe early disease. Despite these limitations, the strengths of this study include careful phenotyping and rich clinical metadata associated with evolving BPD patients and the development of an organotypic *ex vivo* culture system that replicates features of airway disease in preterm infants.

In summary, these results support an early, aberrant airway epithelial cell differentiation deficit that associates with progression to BPD. We propose that this “vimentin-high” airway molecular endotype of BPD may be a driver of the physiological consequences of lower airway obstruction. Leveraging this platform to test for AEC plasticity, maladaptive repair, and extracellular matrix expression may allow for early identification of patients at highest risk for lower airway disease, as well as provide a quantifiable system for screening and testing needed targeted therapies.

**Table 1.**

| **Demographics** | **eBPD** | **Healthy Controls** |
| --- | --- | --- |
| N | 27.0 | 22 |
| sex (% male) | 66.7 | 45.5 |
| age (days for BPD, years for control) | 13.9 +/- 8.9 | 10.3 +/- 4.7 |
| gestational age (weeks) | 24.9 +/-1.9 | term |
| race- white (%) | 70.4 | 63.6 |
| race- Black (%) | 11.1 | 9.1 |
| race- asian | 11.1 | 13.6 |
| race- unknown/not reported or other (%) | 7.4 | 13.6 |
| ethnicity- Hispanic or Latino/a or Latinx (%) | 29.6 | 18.2 |
| ethnicity- Non-Hispanic or Latino/a or Latinx (%) | 63.0 | 81.8 |
| ethnicity- unknown/not reported (%) | 7.4 | 0.0 |
| BW (g) | 716.7 +/- 215.4 | n/a |
| maternal chorioamnionitis (%) | 40.7 | n/a |
| maternal chorioamnionitis unknown (%) | 7.4 | n/a |
| antenatal steroids (%) | 74.1 | n/a |
| postnatal steroids (%) | 96.3 | n/a |
| **Clinical Data** | **eBPD** | **Healthy Controls** |
| FiO2 | 0.36 +/- 0.11 | n/a |
| PEEP | 7.9 +/- 1.8 | n/a |
| jet ventilator (%) | 63.0 | n/a |
| HFOV (%) | 7.4 | n/a |
| ACVG (%) | 25.9 | n/a |
| SIMV (%) | 3.7 | n/a |
| Mild/Moderate BPD (%) | 35.3 | n/a |
| Severe I (%) | 40.7 | n/a |
| Severe Type 2 BPD or death (%) | 37.0 | n/a |

## Supplement

### Detailed Methods

#### Airway Epithelial Cell Organotypic Culture

Unprocessed tracheal aspirates were placed in warm PneumaCult Ex or Ex plus medium, transported to the lab, and then cultured in collagen-coated T25 flasks. These cultures were allowed to proliferate into islands, passaged once more into T25s, then plated at P2 or P3 at 200,000 cells/ 24 well transwell for one week or until confluent.

AECs were then differentiated for three weeks at air liquid interface with PneumaCult-ALI media in the basolateral compartment. In parallel with neonatal AECs, control bronchial AECs from healthy school-age children were collected while underdoing sedated procedures. 4mm Harrell unsheathed bronchoscope cytology brushes (CONMED®) were inserted through an endotracheal tube as we have previously described prior to proliferation of P2 AECs in submerged culture followed by differentiation of P3 AECs at ALI, as previously described (1–4). Quality control was performed in both control and BPD AEC cohorts to ensure they produced an organotypic differentiated epithelial cell culture with mucociliary morphology and without leak indicative of epithelial barrier. TEER measurements were also used on occasion to confirm barrier function.

#### Immunostaining of Organotypic Cultures

AEC cultures were washed with PBS and fixed with 10% formalin for 1h at the room temperature (RT) and labeled by wholemount immunofluorescence staining. After permeation with 0.5% Triton X-100, cells were then blocked with 10% goat serum in PBS for 1h at RT. Incubation with primary antibodies were performed overnight at 4°C, followed by recovery to room temperature and extensive washing with PBS. AECs were incubated with fluorescently labeled stains or secondary antibodies for one hour at room temperature. Thereby, cells were washed with PBS three times before mounting of transwell with DAPI mounting medium on slides. For each AEC sample, two or three locations were imaged. If the cilia stained with R-TUBA distributed evenly on the samples, the images were randomly taken. If the cilia distributed unevenly, the images included both low-density and high-density regions of TUBA+ staining, which were averaged.

The following primary antibodies or stains were used: R-TUBA (Invitrogen, 1:200), R-Vimentin (Invitrogen, 1:100), R-TP63 (Invitrogen,1:100) and Phalloidin (Invitrogen, 1:100). The following secondary antibodies were used: goat anti rabbit Alexa Fluor-488 (Invitrogen, 1:1000) and goat anti rabbit Alexa Fluor-647 (Invitrogen, 1:1000). Phalloidin stain was applied with the incubation of secondary antibodies.

#### Quantitative AEC Differentiation Analysis (High Content Image Analysis)

Leica SP5 and Zeiss LSM 980 confocal microscopes with 10x and 40x oil objective lens was used. For super resolution microscopy, sequential stacks of images were taken along the z-axis at optimal intervals, 30-90 slices were imaged at 0.5 um step size at resolution 1024x1024 pixels. Acquired images were processed using Fiji [19] and Volocity (Quorum technologies) softwares.

Images from Leica SP5 confocal microscope were imported in Fiji/ImageJ2 v2.3.2/1.53q using Bio-format importer. ALI thickness was calculated in ImageJ by counting the number of Z-stacks containing staining and multiplying by the thickness of each layer (0.5 microns). Maximum intensity of every Nuclei Z-stack was projected in order to count total number of cells per image, using manual counting with ImageJ cell counter. Projections of maximum intensity of cilia and actin staining of the apical layer were used as composite images to count the number of apical cells containing cilia using manual counting with ImageJ cell counter. Exported data were analyzed in GraphPad prism v9.4.1.

Cilia per cell analyses included the following procedure for data in Figure 1.

Maximum intensity of Z-stack containing cilia signal was projected and exported in tiff. Those images were then imported in CellProfiler 4.2.1 (5). First illumination was corrected using background correction and gaussian smoothing. Cilia objects were identified from corrected images using Global two classes Otsu thresholding method, with intensity de-clumping. For every image, we reported the total number of objects. Subsequent cilia quantification included measurement of total TUBA+ positive area in 2- 3 areas per transwell; total cell number and ALI thickness were also measured for each sample.

#### RNA Sequencing and Analysis

Bulk RNA sequencing (RNA-seq) was performed on organotypic AEC cultures, with support from the Benaroya Research Institute Genomics and Bioinformatics Cores, with additional details in the supplement. RNA was isolated using RNAqueous™ Total RNA Isolation Kit (Invitrogen) prior to reverse transcription and cDNA generation (SMART-Seq, Takara Bio). We constructed sequencing libraries with the NexteraXT DNA sample preparation kit (Illumina) to generate Illumina-compatible barcoded libraries. Libraries were pooled and quantified using a Qubit® Fluorometer (Life Technologies). Dual-index, single-read sequencing of the pooled libraries was carried out on a HiSeq2500 sequencer (Illumina) with 58-base reads, using HiSeq v4 Cluster and SBS kits (Illumina) with a target depth of 5 million reads per sample. FASTQs were aligned to the human reference genome GRCh38.77 using TopHat (v1.4.1) and gene counts were generated using htseq-count. QC and metrics analysis was performed using the Picard family of tools (v1.134). Low-quality libraries (median CV of coverage > 1, total reads < 5 million) were excluded. Filtered and normalized gene counts were generated from raw counts by trimmed-mean of M values (TMM) normalization and filtered for genes that are expressed with at least 1 count per million total reads in at least 10% of the total number of libraries. Data analysis was performed using the R language using limma (6). Significance in volcano plots was determined by an adjusted p-value cutoff ≤ 0.05 and a log2 fold change ≥ 1 between conditions. Analysis of gene expression programs was performed with Enrichr databases (7–9). Gene set enrichment analysis was also performed using the HALLMARK gene sets (10, 11).

## Supplemental Figure Legends

**Figure E1.**
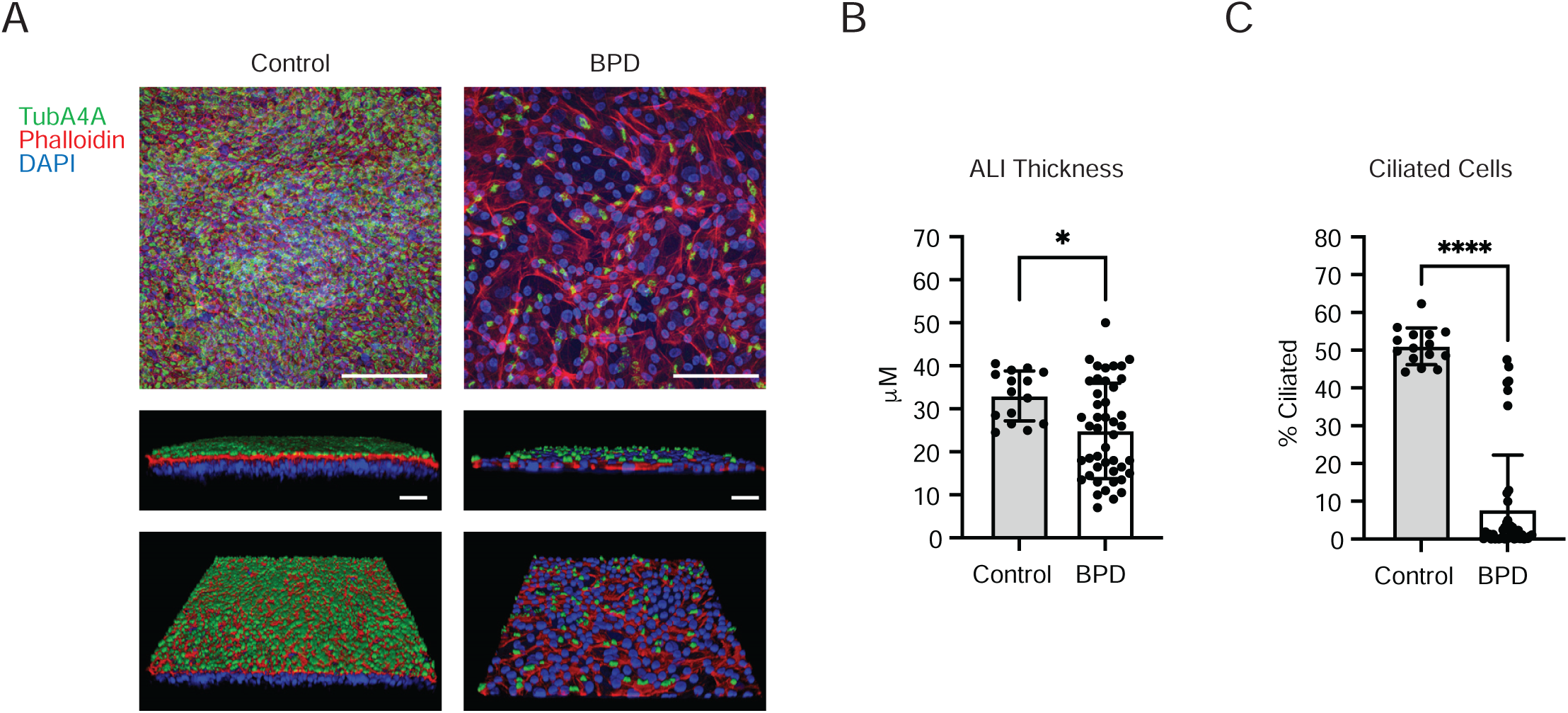
Decreased Apical Cilia and ALI thickness in eBPD Cultures Grown in PneumaCult Ex-Plus Supportive Media. (A) Decreased apical cilia in eBPD cultures identified by immunostaining for TubA4A and actin. (B) Quantification showing decreased ALI thickness and decreased % ciliated cells in organotypic cultures from patients with eBPD (N=6), compared with controls (N=5). One dot is one image, 2-3 confocal images per donor (Green= anti-TubA4A, Red = phalloidin, Blue = DAPI. *p<0.05, ****p<0.0001, Mann Whitney test, magnification bar = 100μm.)

**Figure E2.**
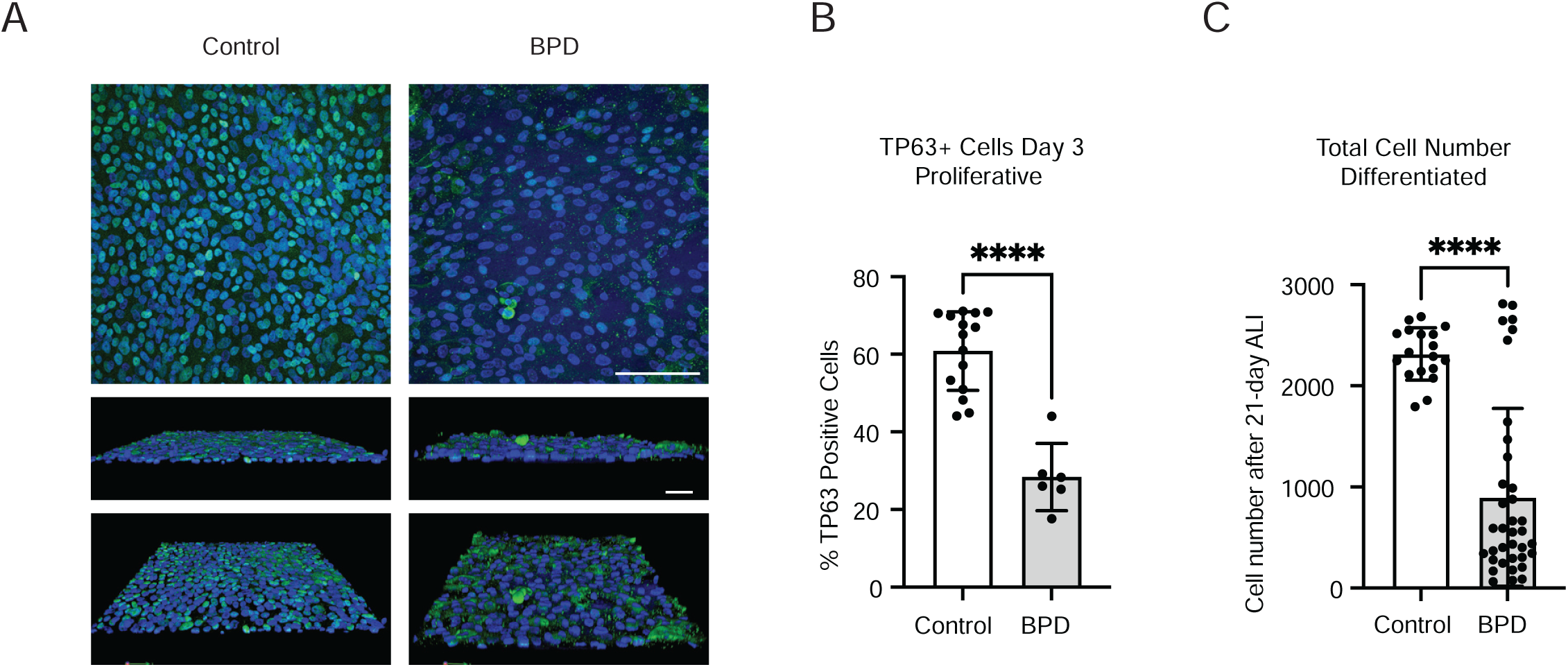
Decreased Proliferative Capacity of eBPD AECs. (A) TP63 immunostaining of proliferative AECs from control and eBPD patients, on day 1 after plating on transwell membranes. Quantification of (B) TP63+ cells in proliferative conditions and (C) total cell numbers in fully differentiated Control and BPD AEC cultures. One dot is one image, with N=2-3 images per condition. Control donor numbers: Day 1=4 donors, Day 3=5, Day 7=6, ALI=9. eBPD donor numbers: Day 1=3 donors, Day 3=3, Day 7=5, ALI=11. (Green= anti-TP63, Blue= DAPI. *p< 0.05, ****p< 0.0001, Mann-Whitney test, magnification bar = 100mm)

**Figure E3.**
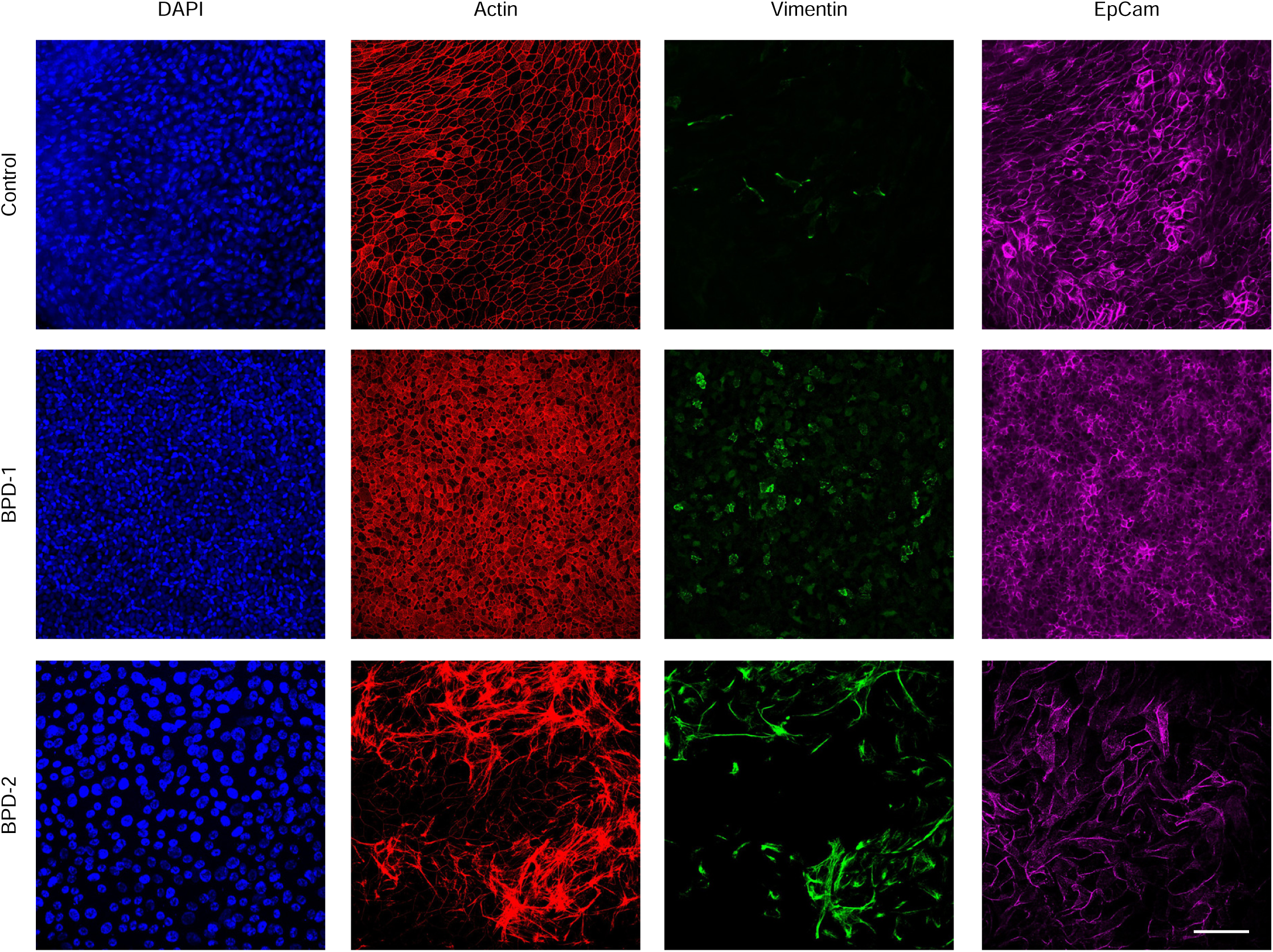
**Increased Vimentin-positive Stromal Cells in Organotypic AEC Cultures from Patients with eBPD. (**Top) Well-differentiated AEC culture from control patient, stained with DAPI, phalloidin, vimentin, and EPCAM antibodies. (Middle) AEC cultures from a patient with eBPD, showing a well-differentiated and polarized culture with increased vimentin and abundant EPCAM staining. (Bottom) AEC from a patient with eBPD, showing a poorly differentiated culture with increased vimentin staining of cells resembling contaminating fibroblasts. (Blue = DAPI, Red = phalloidin, Green = anti-vimentin, purple = anti-EPCAM, magnification bar = 100μm)

